# An important class of intron retention events in human erythroblasts is regulated by cryptic exons proposed to function as splicing decoys

**DOI:** 10.1101/258897

**Authors:** Marilyn Parra, Ben W. Booth, Richard Weiszmann, Brian Yee, Gene W. Yeo, James B. Brown, Susan E. Celniker, John G. Conboy

## Abstract

During terminal erythropoiesis, the splicing machinery in differentiating erythroblasts executes a robust intron retention (IR) program that impacts expression of hundreds of genes. We studied IR mechanisms in the SF3B1 splicing factor gene, which expresses ~50% of its transcripts in late erythroblasts as a nuclear isoform that retains intron 4. RNA-seq analysis of nonsense-mediated decay (NMD)-inhibited cells revealed previously undescribed splice junctions, rare or not detected in normal cells, that connect constitutive exons 4 and 5 to highly conserved cryptic cassette exons within the intron. Minigene splicing reporter assays showed that these cassettes promote IR. Genome-wide analysis of splice junction reads demonstrated that cryptic noncoding cassettes are much more common in large (>1kb) retained introns than they are in small retained introns or in non-retained introns. Functional assays showed that heterologous cassettes can promote retention of intron 4 in the SF3B1 splicing reporter. Although many of these cryptic exons were spliced inefficiently, they exhibited substantial binding of U2AF1 and U2AF2 adjacent to their splice acceptor sites. We propose that these exons function as decoys that engage the intron-terminal splice sites, blocking cross-intron interactions required for excision. Developmental regulation of decoy function underlies a major component of the erythroblast IR program.

## Introduction

Intron retention (IR) is a common variant of alternative splicing in which selected intron(s) are specifically retained in an otherwise spliced and polyadenylated transcript. IR can be developmentally or physiologically regulated as an important component of gene regulation in many cell types (Jacob and Smith 2017). IR transcripts can be stored in the nucleus to be spliced in response to appropriate signals, thus serving as a source of new mRNA (Ninomiya et al. 2011; Boothby et al. 2013; Mauger et al. 2016), or they can represent dead-end RNAs that are degraded in the nucleus (Pendleton et al. 2018). Other IR transcripts are transported to the cytoplasm where they are degraded by nonsense-mediated decay (Wong et al. 2013). Partial diversion of transcriptional output into IR isoforms also functions as a post-transcriptional pathway to modulate expression levels (Wong et al. 2013; Braunschweig et al. 2014; Shalgi et al. 2014; Boutz et al. 2015; Ni et al. 2016). Some IR transcripts can be recruited to ribosomes to function in translation (Li et al. 2016), while other might serve as miRNA sponges in the nucleus (Schmitz et al. 2017).

The diversity of IR programs and functions suggests that a number of regulatory pathways exist, allowing cells to integrate multiple inputs in order to independently regulate different subsets of IR transcripts. Consistent with this notion, coherent subsets of genes can be regulated by IR, especially those encoding RNA binding proteins and spliceosomal factors (Shalgi et al. 2013; Wong et al. 2013; Braunschweig et al. 2014; Boutz et al. 2015; Pimentel et al. 2016). How these sub-programs are regulated is not well understood. Recent studies have implicated transcription rate/pausing (Braunschweig et al. 2014), specific splicing factors (SRSF4 and HNRNPLL), and DNA and protein methylation factors (by MECP2, in granulocytes, and by PRMT5, in glioblastoma) as important effectors of IR (Cho et al. 2014; Boutz et al. 2015; Braun et al. 2017; Wong et al. 2017). In the brain, signaling-dependent splicing was observed to require NMDA-type glutamate receptor or calmodulin-dependent protein kinase pathways for removal of retained introns upon neuronal activation (Mauger et al. 2016). There are also a few recent examples showing that individual IR events can be regulated by feedback mechanisms to control physiological pathways (Park et al. 2017; Pendleton et al. 2017; Pirnie et al. 2017).

Terminal erythropoiesis is an excellent model system for studies of IR. Primary erythroblasts differentiating in culture carry out a robust IR program (Edwards et al. 2016; Pimentel et al. 2016) impacting the expression of many important erythroid proteins. In mature erythroblasts, IR transcripts comprise 25-50% of steady state RNA for genes encoding essential splicing factors (SF3B1 and others), mitochondrial iron importers required for heme biosynthesis (mitoferrins, encoded by SLC25A37 and SLC25A28), and major cytoskeletal proteins (alpha spectrin, encoded by SPTA1). Moreover, the cellular complement of IR events is continuously remodeled in a differentiation stage-dependent manner through the combined effects of differentiation-independent and –dependent IR networks (Pimentel et al. 2016).

In the current study, we focused on regulatory mechanisms for a subset of genes represented by SF3B1. SF3B1 is an essential splicing factor that functions in normal 3’ splice site regulation. Mutations in the SF3B1 gene are found in many MDS (myelodysplastic syndrome) patients, where they induce RNA processing errors due to altered 3’ splice site selection (Obeng et al. 2016) and changes in exon skipping (Jin et al. 2017). We used a combination of comparative genomics, RNA-seq and RT-PCR analysis, and minigene splicing reporters to discover highly conserved decoy exons that strongly influence the level of intron 4 retention. Furthermore, bioinformatics analysis of RNA-seq data demonstrated that decoy exons are a common feature of large (>1kb) retained introns. We propose that decoy exons interact nonproductively with intron-terminal splice sites to block intron excision, and that this mechanism regulates a critical subset of IR events in differentiating erythroblasts.

## Results

### SF3B1 retained intron 4 (i4) harbors cryptic exon(s) that are highly conserved

Comparative genomic analysis showed that SF3B1 i4 is extremely conserved among vertebrate genomes (Figure 1A). Three 125-200nt regions are 93-98% identical from chicken to human, and core areas of two are 79-94% identical from zebrafish to man (Figure S1). Sequence inspection revealed several pairs of consensus 3’ and 5’ splice sites in these ultra conserved regions, as well as in three additional conserved regions, predicting six short exons of 29-56nt (Figure 1A, E4a-E4f). The extraordinary conservation of these exons to fish (E4d and E4e), reptiles (E4b, E4c, and E4f), and mammals (E4a) is shown in Figure S1, and suggests that these cryptic exons might have important function(s) in SF3B1 regulation. Consistent with the idea that the splicing machinery might recognize these exons, we discovered that the 3’ splice site factors U2AF1 and U2AF2 cross-linked to five of the six exons in K562 (erythroleukemia cells) eCLIP experiments (Figure 1a, lower panels) and to all six exons in HEPG2 cells (not shown). Since all except E4c would introduce premature termination codons (PTCs), we hypothesized a non-coding function for these exons.

**Figure 1.**
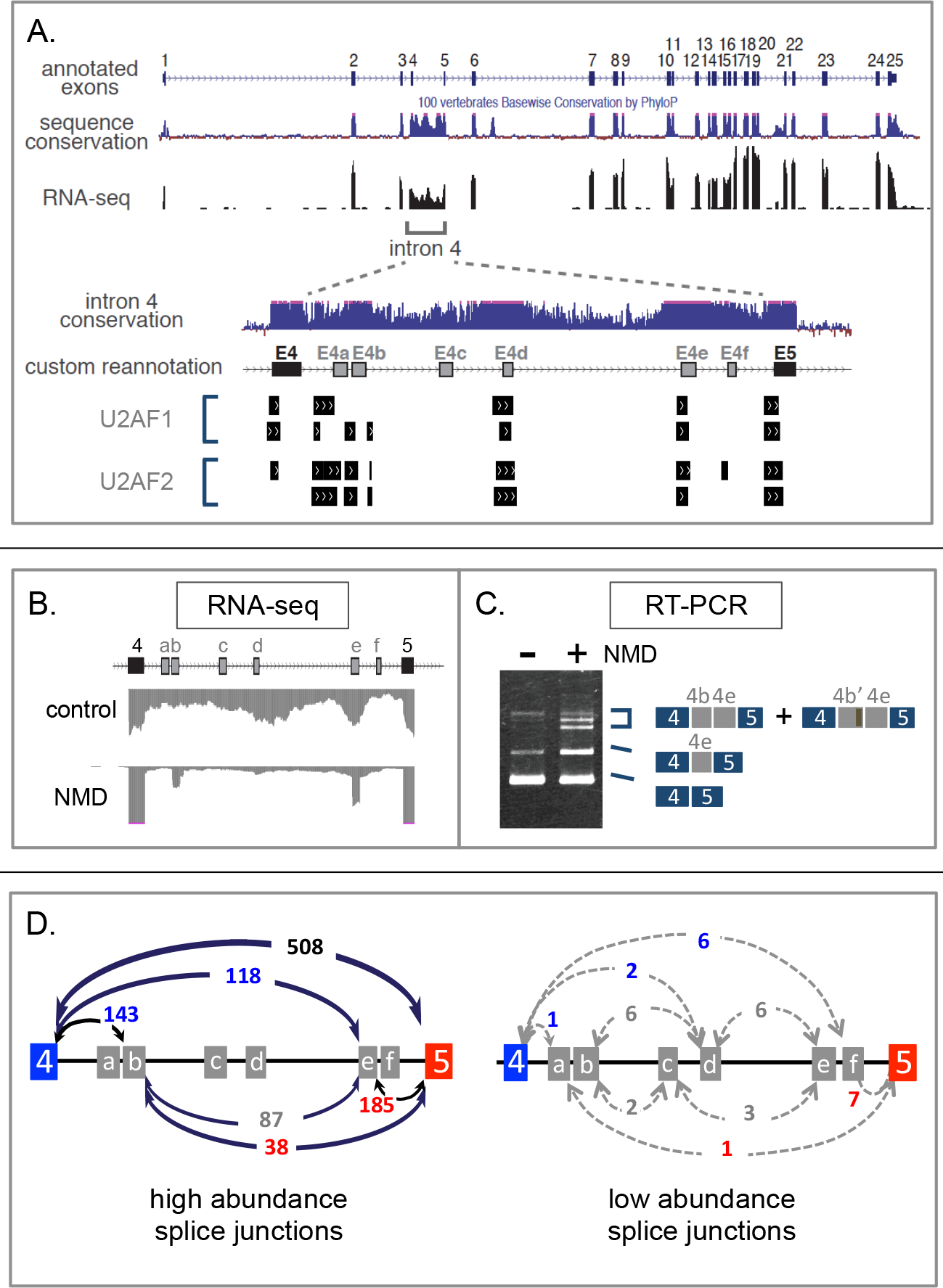
Structure and expression of SF3B1 intron 4. A. Top portion, genome browser tracks showing exons, conservation, and RNA-seq coverage of the SF3B1 gene. Bottom portion, close-up of i4 region Showing predicted cryptic exons E4a-E4f and binding data for U2AF subunits. B. RNA-seq coverage in control cells vs NMD-inhibited cells. C. RT-PCR analysis of cryptic exon expression. D. plots showing RNR-seq junctions in NMD-inhibited D16 erythroblasts.

In order to study expression of cryptic exons in SF3B1 intron 4, we performed transcriptome analysis of early erythroblast progenitors (culture day 9, D9) and mature erythroblasts (culture day 16, D16) that had been treated with inhibitors of nonsense-mediated decay (NMD). This strategy increased the relative abundance of transcripts containing premature termination codons (PTC), allowing us to validate expression of two cryptic exons (E4b and E4e) in the RNA-seq profiles and by RT-PCR (Figures 1B-C). Furthermore, analysis of individual RNA-seq reads revealed new splice junctions that validate expression of all six predicted cassettes, and indicate extensive connections between and among these cryptic exons and the flanking constitutive exons 4 and 5 (Figure 1D). Two important features were noted in this analysis. First, each cryptic exon was represented by at least one unique RNA-seq read that connected it to both upstream and downstream exons, confirming its ability to be recognized and spliced as a discrete exon. Second, splice junctions that link constitutive exons 4 and 5 with intron-internal sites were surprisingly abundant. In the D16 sample, E4-E4b and E4-E4e junctions together represented ~33% of splice junction reads (261/778) that connect E4 to downstream sequences, while ~30% of E5 upstream splice junction reads (223/739) involved E4b or E4e. Splice junctions that connect E4b to E4e were also common. At lower frequency, splice junctions connected the other cryptic exons to each other and to the flanking constitutive exons (Figure 1D).

Together these experiments confirmed that the intron-terminal splice sites of i4 interact with internal splice sites associated with cryptic exons. We hypothesized that the cryptic exons might function as decoys whose splice sites could compete with the cross-intron interactions required for intron excision, and thereby might promote intron retention. We reasoned that many of these interactions might exhibit “leaky” splicing, leaving behind splice junctions that serve as indirect evidence for these interactions. This model suggested several testable predictions: that deleting decoy exons or mutating their splice sites should reduce intron retention; that known 3’ splice site factors should cross-link to the cryptic junctions; and that analogous decoy exons / splice junctions should be common in retained introns.

### Decoy exons promote SF3B4 i4 retention

We investigated the sequence requirements for intron retention using a series of minigene splicing reporters. The wild type (WT) construct contains a 4.7kb fragment of the SF3B1 gene spanning exons 3-6 and includes full-length intron sequences in this region (Figure 2A). This reporter was spliced to produce two major products when transfected into K562 cells: a fully spliced RNA containing exons 3-6, and a larger transcript that has excised introns 3 and 5 but specifically retained i4 (Figure 2B, left lane). In contrast, little intron retention was observed in HEK (human embryonic kidney) cells (Figure 2B, right lane), demonstrating cell type-specific regulation of i4 retention.

**Figure 2.**
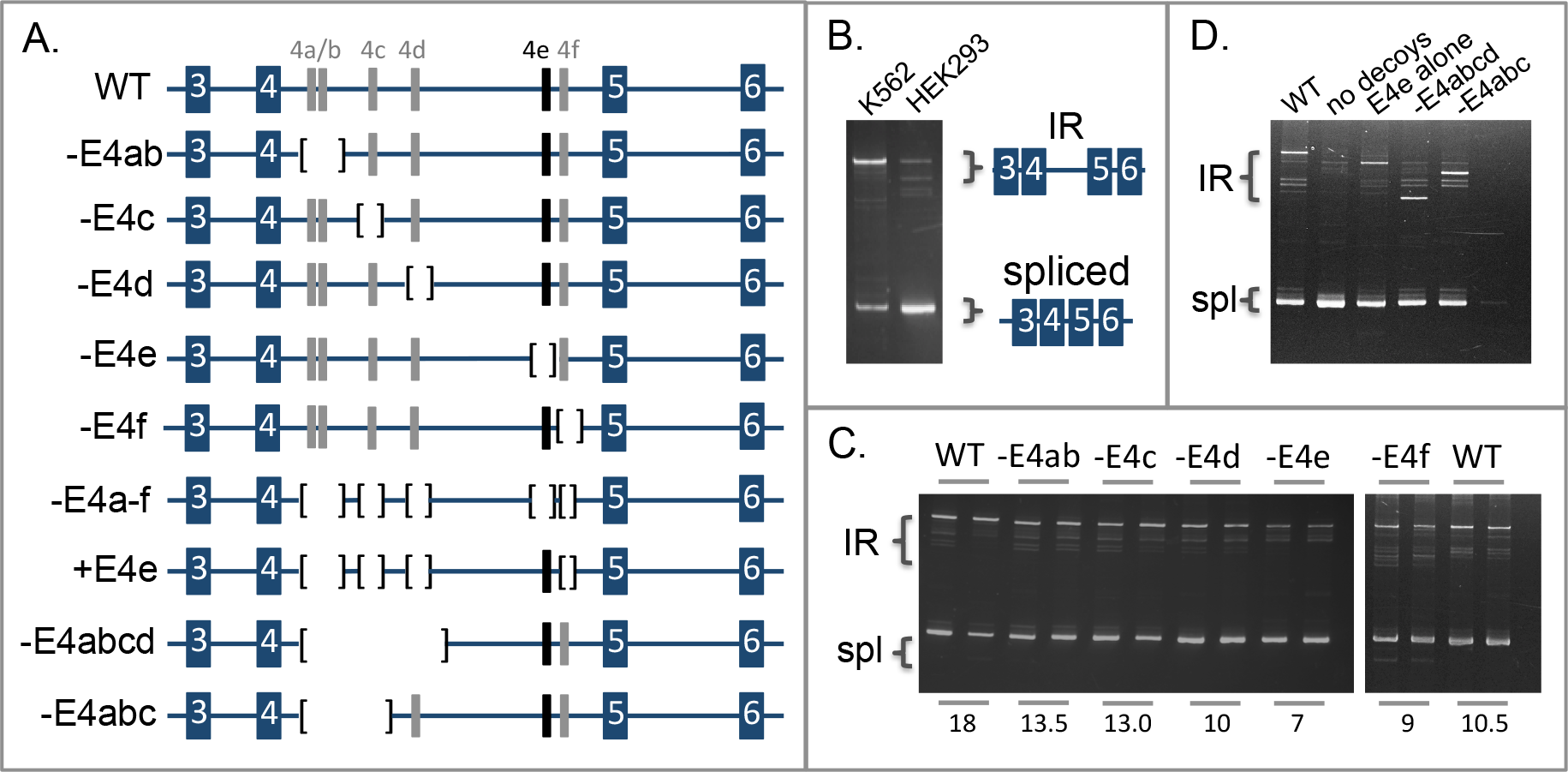
Intron retention assays in minigene splicing reporters. A. Structure of minigene splicing reporters spanning the E3-E6 region of SF3B1. Shown are the wild type reporter (WT) and a series of variants with deletions of candidate decoy exons. B. Splicing assay performed with the wild type reporter in K562 and HEK293 cells. C. Splicing assays show that single-decoy deletions have mild effects on IR. Numbers below the lanes indicate apparent IR percent as determined by densitometry, corrected for differences in band size. D. Splicing assays with multi-decoy deletions show that even a single decoy (E4e) can promote IR.

To test whether the cryptic exons in intron 4 function as decoys to promote intron retention, we constructed a series of splicing reporters in which these putative decoys were deleted. Deletion of individual exons resulted in slightly reduced IR compared with the WT construct, with E4e having the greatest effect (Figure 2C). Deletion of all six decoys resulted in a smaller intron with greatly reduced IR (Figure 2D, compare first two lanes), while adding back a single copy of E4e restored substantial retention (3^rd^ lane). As a control, we showed that loss of IR in the decoy-deficient reporter was not due to the smaller size of the intron, since E4e-containing constructs with similar-sized introns retained significant IR (last two lanes).

### Decoy exon E4e can promote IR at heterologous sites

We next investigated whether E4e could promote IR in different intronic contexts. First, the ability of E4e to function at a heterologous position in i4 was tested. Indeed, the reduction in IR observed when E4e was deleted from the wild type construct (Figure 3B, compare WT with Δ4e) was rescued when E4e was substituted at the E4d site (mut21).

**Figure 3.**
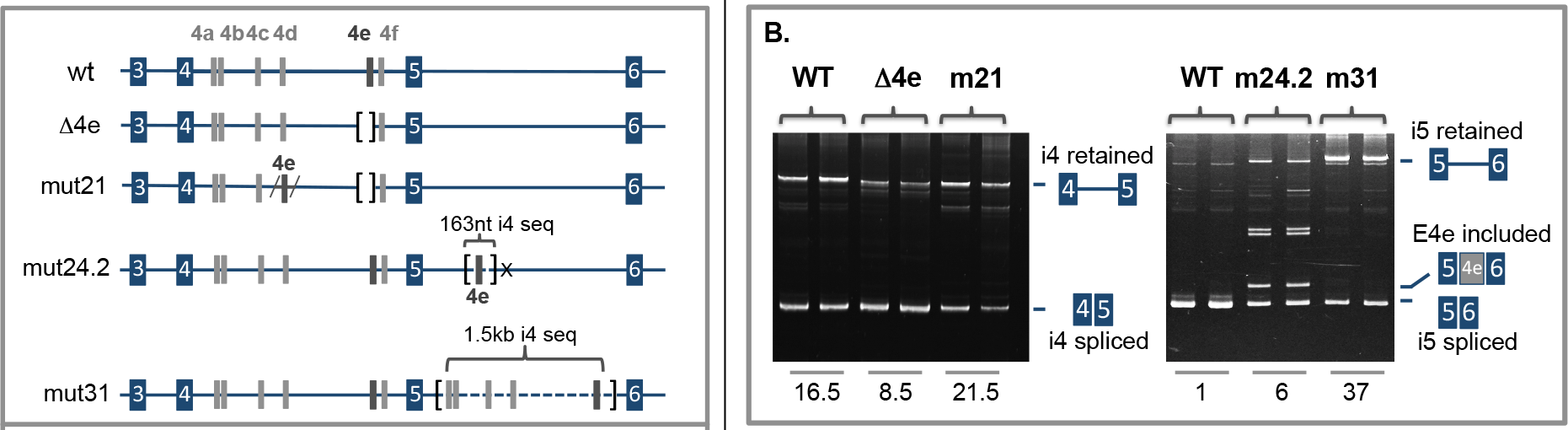
Testing IR activity of a decoy exon at heterologous sites. A. Structure of minigene splicing reporters testing function of E4e in a different region of i4 (mut21) or in i5 (mut24.2, mut31). B. RT-PCR analysis of intron retention showing that decoy exons can promote retention at heterologous sites in the same intron or in another intron. Numbers below the lanes indicate apparent IR percent as determined by densitometry, corrected for differences in band size.

We then asked whether IR-promoting elements in i4 could induce IR in a non-retained intron. Intron 5 (i5) in the SF3B1 gene ordinarily exhibits negligible IR in either the endogenous gene (not shown), or in splicing reporters (Figure 3B, right gel, lanes WT). In construct mut31, most of i5 (~1.1kb) was replaced with a similar length of i4 including E4a through E4e, but ~160nt of the natural i5 sequence was maintained at either end. This arrangement induced substantial retention of i5. When E4e alone was inserted at a random site within i5 (mut24.2), modest IR activity was observed, but it was accompanied by substantial inclusion of E4e. Thus, decoy exons can promote IR in a heterologous intron, independent of specific sequences at the intron-terminal splice sites or flanking constitutive exons. The neighboring sequence environment of a decoy is critical, however, since a “permissive” sequence context in intron 5 apparently enhanced exon E4e inclusion rather than the IR phenotype observed in intron 4. In fact, for this experiment it was necessary to mutate a cryptic 5’ splice site in intron 5 (marked by the “X”), because otherwise the major product was strong inclusion of an aberrant E4e-related exon.

### Candidate decoy exons occur in many retained introns

Studies of IR-promoting elements in introns of OGT (Park et al. 2017) and ARGLU1 (Pirnie et al. 2017) previously showed that internal intronic elements, including at least one cryptic cassette exon, can promote IR. In the OGT gene, an intron splicing silencer (ISS) inhibits intron 4 removal, thus favoring intron retention and contributing to O-GlcNAc homeostasis (Park et al. 2017). This OGT intron was strongly retained in erythroblasts, and splice junction data revealed a novel ~143-153nt cassette exon that overlaps the ISS and possesses alternative 3’ and 5’ splice sites (Figures S2 and S3). In ARGLU1, a highly conserved cassette exon was previously shown to promote IR in HeLa cells (Pirnie et al. 2017). This exon was expressed in erythroblasts as a 58-200nt cassette, depending on alternative splice site usage (Figures S2 and S3). Further inspection of splice junction reads revealed candidate decoy cassettes in retained introns identified earlier in DDX39B, SPTA1, KEL, SNRNP70, and FUS transcripts ((Edwards et al. 2016; Pimentel et al. 2016); Figure S3). In each case, the putative decoy is flanked on both sides by retained intron(s). This genomic configuration is consistent with our earlier finding that PTC-containing alternative exons are often situated between two consecutive retained introns (Pimentel et al. 2016). Together these observations suggested that decoy cassettes might be a common phenomenon among retained introns.

To address the genome-wide potential for decoy exon-mediated IR, we correlated IR and splice junction data for all introns in erythroid-expressed genes. Among 20,534 retained introns identified in the early erythroblast D9 sample, 770 predicted cassettes were identified (Supplemental Table 1). Late erythroblast D16 cells, that have down-regulated expression of many genes, exhibited 411 cassettes in 9677 retained introns. Interestingly, the incidence of candidate decoys was strongly dependent on intron length; both D9 and D16 transcriptomes exhibited a much higher frequency of cassettes in longer retained introns (≥ 1kb; 12-17%) than in shorter retained introns (~2%). Consistent with this observation, the median length of retained introns with cassettes was much greater (1176-1280nt) than that of retained introns lacking cassettes (258-311nt). These results provided the first clue that distinct IR mechanisms might preferentially regulate introns of different lengths.

**Table 1.**

| sample | R-introns>1kb |  | % Cassette | R-introns<1kb |  | % Cassette |
| --- | --- | --- | --- | --- | --- | --- |
|  | total | +cass. |  | total | +cass. |  |
| D9 | 4038 | 482 | 11.9 | 16,496 | 288 | 1.7 |
| D16 | 1436 | 238 | 16.6 | 8241 | 173 | 2.1 |
| sample | N-introns>1kb |  | % Cassette | N-introns<1kb |  | % Cassette |
|  | total | +cass. |  | total | +cass. |  |
| D9 | 82,853 | 3600 | 4.3 | 36,664 | 179 | 0.5 |
| D16 | 78,454 | 3443 | 4.4 | 41,360 | 244 | 0.6 |

However, the analysis above does not take into account that some of cassettes are likely to represent “conventional” NMD-inducing exons that play no role in IR. We reasoned that the real frequency of IR-promoting cassettes could be estimated by comparing cassette frequency in retained vs non-retained introns, since the latter group by definition lacks IR-promoting cassettes. Table I shows that the frequency of cassettes was 3-4-fold higher in the large retained introns (R-introns) than in the comparable non-retained introns (N-introns), for both D9 and D16 transcriptomes. Based on the larger overall numbers of N-introns, and the greater frequency of cassettes in R-introns, we conclude (1) that most noncoding cassette exons in erythroblasts are located in non-retained introns and are not relevant to IR, and (2) that a majority of cassettes located in R-introns likely do act to promote IR.

These observations support the concept of two mechanistically distinct classes of retained introns: longer introns in which decoy exons often promote IR, and shorter introns that are retained via decoy-independent mechanism(s). Overall, the data indicates that several hundred introns may be regulated by the decoy mechanism in human erythroblasts. The true number could be higher, if some functional decoys were not detected because they are poor spliced and did not generate splice junction reads.

### Heterologous decoy exons can promote intron 4 retention

We proposed that heterologous IR-associated cassette exons could substitute for SF3B1 E4e to promote retention of i4. To explore this hypothesis, we assayed their function in minigene splicing reporters (Figure 4A). Each candidate decoy was inserted into reporter ΔE4e, which has low intrinsic IR, and changes in retention were assessed in transfected K562 cells (Figure 4B). As a positive control we tested the 58nt ARGLU1 exon, and in parallel tested several additional candidate decoys. In the minigene assay, the ARGLU1 decoy exhibited strong IR activity as indicated by increased intensity of the IR bands and decreased intensity of the spliced products compared with ΔE4e (Figure 4B, compare lanes 2 and 3). Even stronger IR activity was associated with the 146nt OGT cassette (lane 4). DDX39B intron 6 was predicted to encode a decoy of 122-369nt, depending on use of alternative splice sites; this also decoy exhibited strong IR activity (lane 5). A 60nt cassette in SNRNP70 intron 7 showed variable but relatively low activity (lane 6). In contrast, neither an 85nt predicted cassette in FUS intron 7, nor a 33nt cassette in KEL intron 6, had detectable IR activity (lanes 7-8).

**Figure 4.**
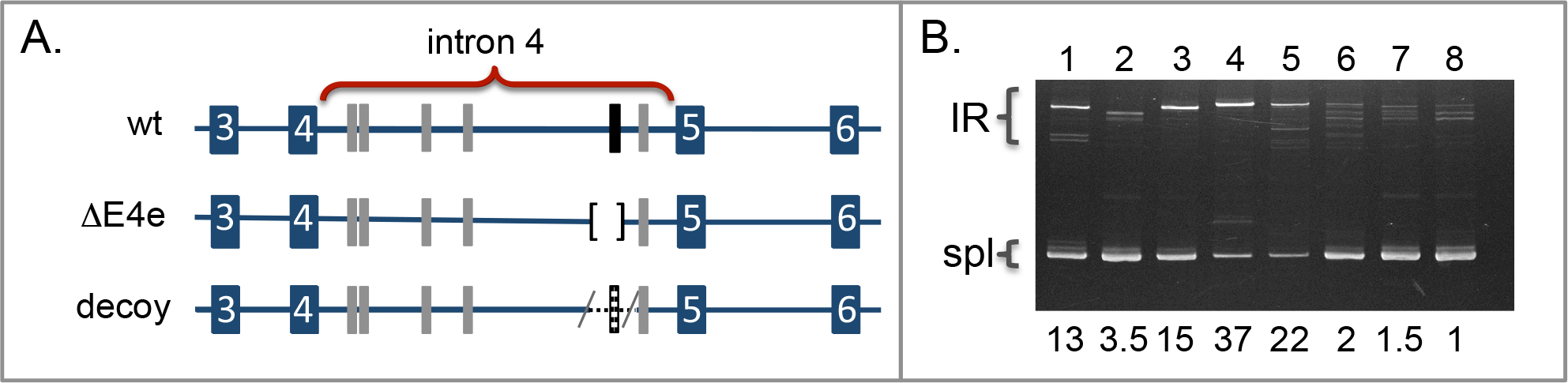
A. Splicing reporters used to assay heterologous decoys. Construct “decoy” had E4e deleted and replaced with candidate decoys from other genes. B. RT-PCR analysis of intron retention activity associated with heterologous decoys in K562 cells. Lane 1, wild type; lane 2, ΔE4e; lanes 3-8, replacement of E4e with candidate decoys from ARGLU1 (lane 3); OGT (lane 4); DDX39B (lane 5); SNRNP70 (lane 6); FUS (lane 7); and KEL (lane 8). IR, intron retention product; spl, spliced product. Numbers below the lanes indicate apparent IR percent as determined by densitometry, corrected for differences in band size.

### Decoy splice sites are critical for IR function

Using the strong OGT decoy exon as a model, we tested the prediction that decoy splice sites play a critical role in promoting IR. The decoy-deficient SF3B1 splicing reporter exhibited little or no retention (Figure 5, lanes 1-2), while insertion of the OGT decoy promoted strong IR (lanes 3-4). Mutating GT dinucleotides to CT at both 5’ splice sites of the decoy exon (Figure S2) essentially abrogated IR in favor of completely spliced transcripts (Figure 5, lanes 5-6). At the 3’ splice site, mutation of the two alternative AG dinucleotides also eliminated retention of the full-length intron (Figure 5, lanes 7-8), but had a more complex phenotype due to activation of a cryptic 3’ splice site that decreased the size of the decoy exon. Fully spliced (intron excision) transcripts were not observed in this mutant. Instead, (truncated) decoy inclusion products were generated together with partial IR products retaining only the downstream intron. The fact that splice site mutations almost completely abrogated retention of the full intron strongly supports the decoy exon model.

**Figure 5.**
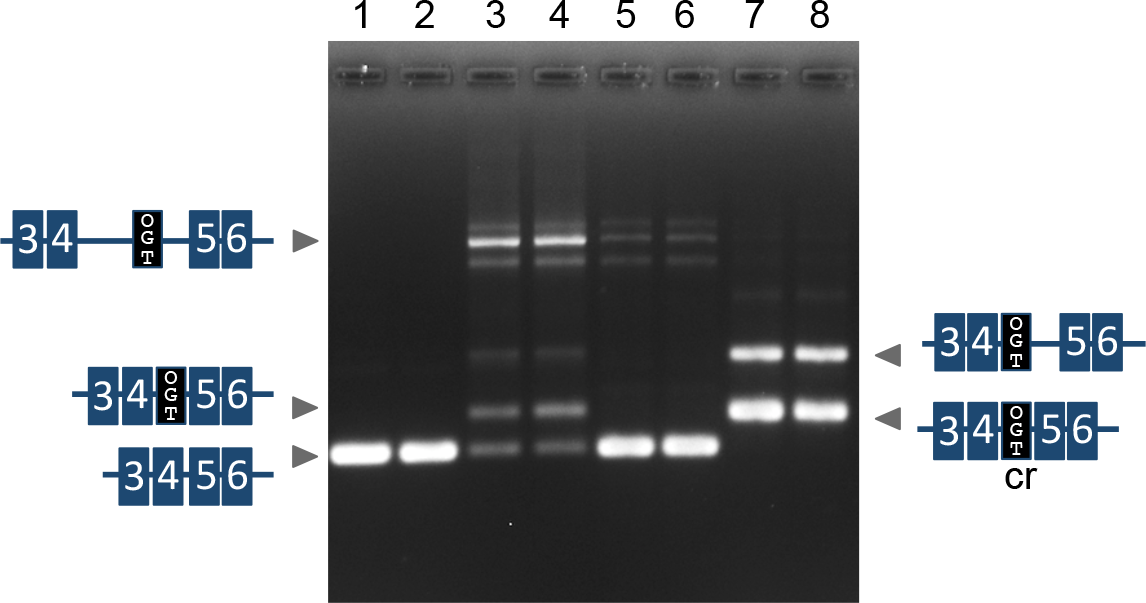
Decoy splice site mutations affect the balance of splicing outcomes involving a retained intron. Gel shows splicing results from the following reporters: lanes 1-2, SF3B1 splicing reporter -E4a-E4f (no decoys); lanes 3-4, OGT decoy; lanes 5-6, OGT decoy with mutated 5’ splice sites; lanes 7-8, OGT decoy with mutated 3’ splice sites. 5’ splice site: WT= ATGgtaacgggt; mut5’= ATG<u>c</u>taacgg<u>c</u>t; 3’ splice site: WT=ttta<u>g</u>aa<u>g</u>GTT; mut3’=ttta<u>c</u>aa<u>c</u>GTT. Underlined nucleotides were altered from “g” in WT to “c” in mutants. WT, wild type; cr, cryptic splice site.

### Binding of U2AF1 and U2AF2 to decoy exons

If decoy exons compete with intron terminal splice sites to enhance IR, the decoy model predicts that decoy splice junctions must be recognized by the splicing machinery. In individual cases, we already showed (Figures 1 and S3) that U2AF1 and 2 binding sites align with the SF3B1, OGT, ARGLU1, and DDX39B decoy exons, but little or no binding was associated with candidate decoys that did not promote IR. Here we tested the association between decoy exons and U2AF binding more globally by examining ENCODE eCLIP data that define U2AF1 and U2AF2 binding sites in the K562 transcriptome. Good binding peaks for both U2AF1 and U2AF2 were observed at the expected location upstream of these exons (Figure 6), similar to binding patterns for a control set of known cassettes expressed in K562 cells (native cassette exons). This binding profile supports the model that many decoy exon indeed bind to U2AF splicing factors as predicted by the decoy model.

**Figure 6.**
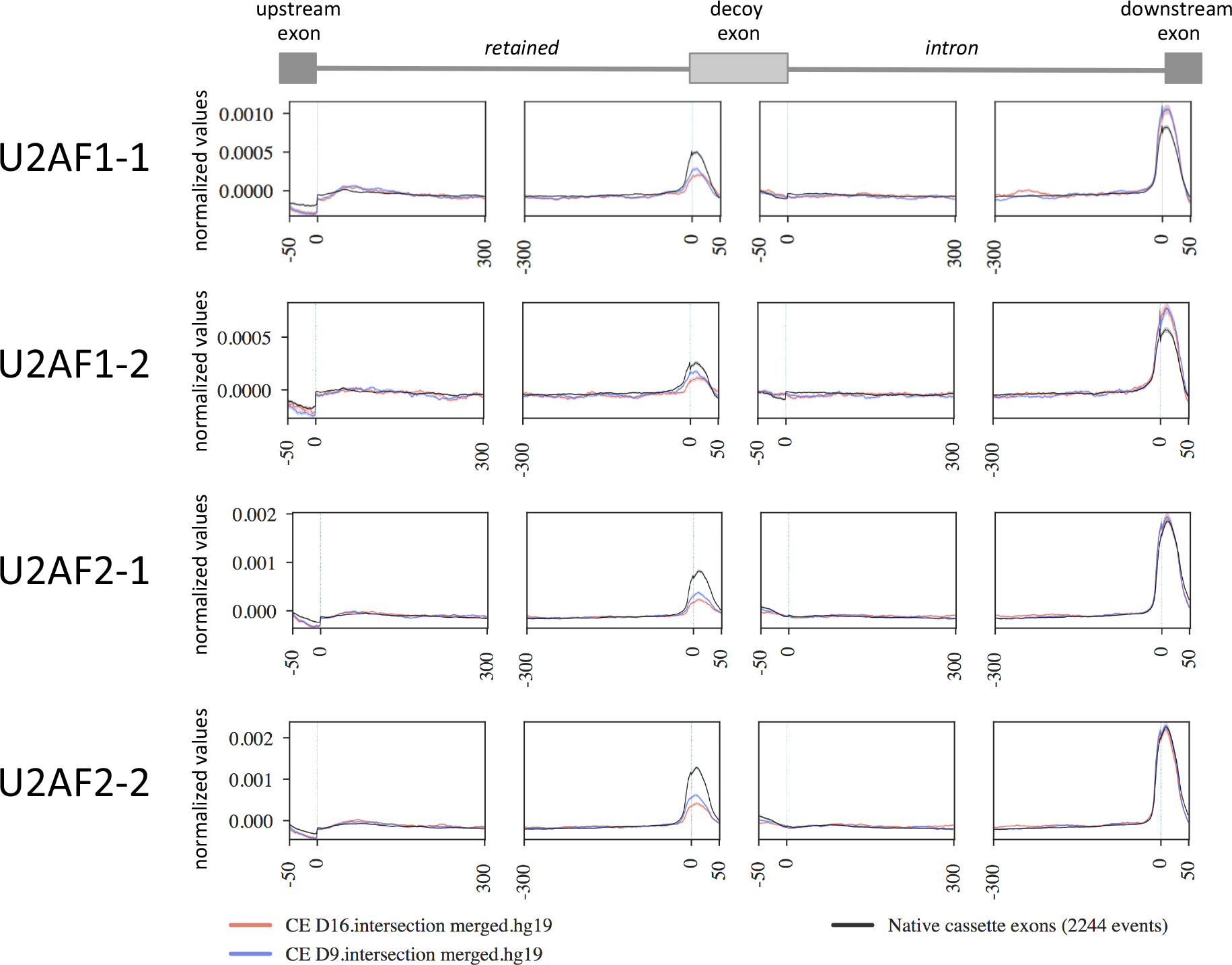
U2AF binding profiles in retained introns with candidate decoys. Enhanced CLIP binding profiles for U2AF1 and U2AF2 in K562 cells. Profiles show enriched binding for U2AF factors near candidate decoys was similar to that observed for a control set cassette exons in the same cells.

## Discussion

Experimental and computational data support the concept that decoy-mediated IR is a novel mechanism for regulating an important component of the erythroid IR program. Experimental analysis of SF3B1 minigene splicing reporters demonstrated that highly conserved decoy exons are required for optimum intron retention, that decoy exons can induce IR in an ordinarily non-retained intron, and that heterologous decoys from other retained introns can promote SF3B1 i4 retention. In parallel, computational analysis of RNA-seq data from NMD-inhibited cells suggested that a broader program of decoy exon-regulated IR is executed during terminal erythropoiesis. Our model expands the range of splicing outcomes available to noncoding cassette exons. As shown in Figure 7, skipping of these exons leads to productive splicing and generation of translatable mRNAs; inclusion of decoy exons yields unstable transcripts that are subject to NMD; and nonproductive interaction with intron-terminal splice sites represents a novel IR outcome. Notably, the decoy exon mechanism greatly extends the model reported recently for ARGLU1, where it was proposed that unproductive splicing complexes assembled at the alternative exon disfavor intron splicing so as to promote its retention (Pirnie et al. 2017). The model is also consistent with previous reports that RIs are enriched adjacent to alternative exons (Braunschweig et al. 2014; Pimentel et al. 2016). Depending on variables including splice site strength, nearby enhancer and silencer elements, and physiological context, we envision that dual-function cassettes may post-transcriptionally direct transcript outcomes preferentially towards NMD or towards IR. At the extremes, some cryptic cassettes may function solely to induce NMD, while others may function predominantly to promote IR. The latter subset of decoys might be difficult to detect using splice junction criteria, since they would rarely splice to the flanking constitutive exons.

**Figure 7.**
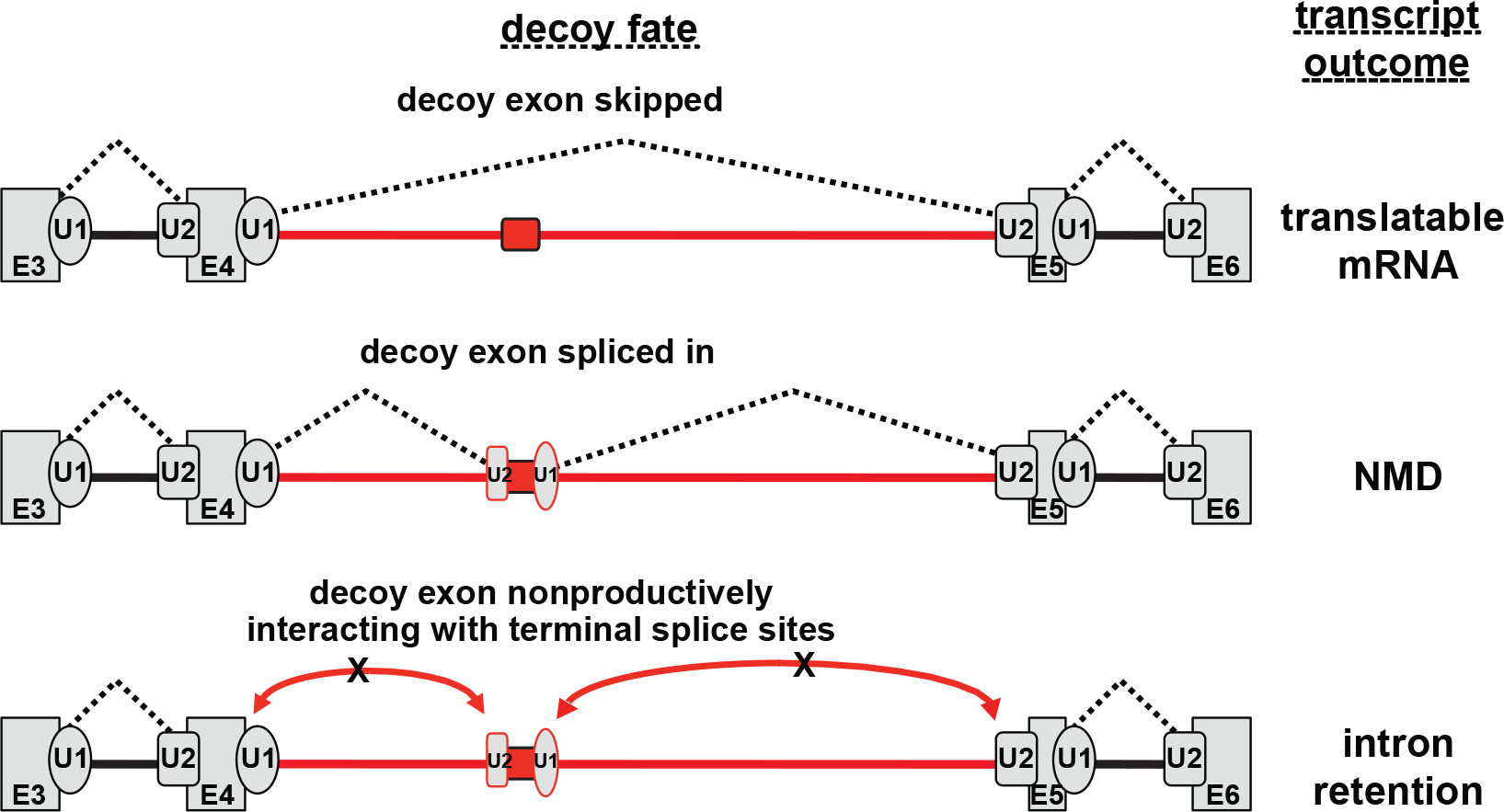
Decoy exon splicing model. Behavior of the decoy exon (red) can dictate three distinct fates for the pre-mRNA: skipping of the exon leads to production of mRNA; inclusion of the decoy generates an NMD-sensitive isoform; and nonproductive interaction with flanking exons yields the intron retention transcript. Dotted lines indicate splicing; red curved lines indicate nonproductive interactions (no splicing). Decoy function could be regulated through the action of nearby enhancer and/or silencer elements (not shown).

The role of decoy-mediated IR in terminal erythropoiesis remains to be investigated. In human erythroblasts, our data indicate that decoy-regulated RIs comprise several hundred erythroblast retention events, mostly involving >1kb introns that are mechanistically distinct from the more numerous population of smaller RIs that generally lack decoys. Given our earlier finding that several major spliceosomal factors possess highly retained introns (Pimentel et al. 2016), and new data showing that some of these possess IR-promoting decoys, we suggest this mechanism could modulate changes in splicing capacity of late stage erythroblasts as they reduce gene expression in preparation for enucleation. Another possibility is that regulated IR could contribute to balanced expression of competing or cooperating genes. For example, modulation of IR plays an important role in regulating expression for competing OGT and OGA enzymes to ensure O-GlcNAc homeostasis (Park et al. 2017), and the documented decoy exon in OGT likely contributes to that control. Speculatively, since the alpha spectrin gene SPTA1 has a decoy exon in retained intron 20, IR could help ensure balanced expression of alpha and beta spectrin subunits, two structural polypeptides that form long heterodimers and provide mechanical support to the red cell membrane skeleton.

The molecular mechanism by which decoy sites can non-productively engage intron-terminal splice sites requires further study. Decoy splice site function must be carefully tuned, since strong splice sites would favor decoy exon inclusion over intron retention, while splice site-inactivating mutations can abrogate IR-promoting activity. One feature that may be critical is the presence of multiple competing splice sites at the decoy exon. Multiple 5’ and/or 3’ splice sites are a feature of the decoy exon in ARGLU1 (Pirnie et al. 2017) as well as OGT and DDX39B (Figure S2). Moreover, even when explicit splice junction multiplicity is not evident, some decoys possess potentially competing splice site motifs in the proximal introns (results not shown). On the other hand, cassette exons encoded in retained introns from FUS and KEL did not appear to have alternative splice sites, and did not exhibit IR activity in the splicing reporter assay.

Competition between splice sites is a fundamental governing principle of alternative splicing. The concept that competing sites can be spliced inefficiently, functioning mainly as decoys to block functional use of other sites, is also well grounded in previous work. An early example was the Drosophila P-element gene, where U1 snRNP binding to an exonic pseudo-5’ splice site was shown to compete with the normal downstream 5’ splice site to promote retention of the adjacent intron in somatic cells (Siebel et al. 1992). Subsequent studies showed that a decoy 3’ acceptor can engage the 5’ splice site of an upstream exon to promote its skipping in caspase-2 (Cote et al. 2001) and other transcripts (Havlioglu et al. 2007). The concept that a noncoding exon might engage splice sites of both flanking exons to induce IR was suggested recently (Pirnie et al. 2017), and the decoy exon data presented here greatly expands the potential role of decoy splice sites in regulation of splicing outcomes. Interestingly, the appreciation of decoy exons might help to explain recent observations in other systems. Far-distal branchpoints located >100nt upstream of annotated 3’ splice sites are more common in retained introns than in constitutive introns (Pineda and Bradley 2018); the increased frequency in retained introns could be due in part to branchpoints associated with cryptic decoy exons rather than the constitutive exons. In other studies, U2AF was shown to bind at numerous intronic locations not corresponding to known 3’ splice sites (Shao et al. 2014). Our data suggest that some of these likely correspond to decoy exons that promote intron retention.

In conclusion, the decoy model represents a new and distinctive component of the erythroid alternative splicing program. We speculate that developmentally regulated IR events in other cell types may also be regulated by decoy exons, and that physiological control of such programs could be mediated by context-dependent combinations of splicing enhancer and silencer proteins that impact recognition of the decoys. We anticipate that analogous subsets of IR events might be regulated by decoy mechanisms in other developmental or physiological contexts. Possible candidates could include differentiating granulocytes (Wong et al. 2013; Wong et al. 2017), activated T cells (Ni et al. 2016), stimulated neurons (Mauger et al. 2016), cells subjected to proteotoxic stress (Shalgi et al. 2014), proliferating vs differentiated muscle cells (Llorian et al. 2016), differentiating germ cells (Naro et al. 2017), etc. Identifying the RBPs that regulate these decoy-dependent programs will be an important goal of future studies in this area.

## Methods

### Erythroblast culture

Cells were cultured as described previously (Hu et al. 2013). For NMD experiments, cultures were divided and one half of each culture was incubated with 100ug/ml emetine for 8 hours and 100ug/ml cycloheximide for 4 hours. Experiments were done in biological duplicates for a total of eight samples (two differentiation stages, two conditions plus or minus NMD inhibitors, and two biological replicates.

### RNA and RNA library Preparation

RNA was isolated from erythroblasts (4x10^6^ cells from day 9 and 16x10^6^ cells on day 16) according to the manufacturer’s instructions (Qiagen). Sequencing libraries were prepared from 500 ng of total RNA using the NEBNext Poly(A) mRNA Magnetic Isolation Module (catalog number E7490, protocol revision 5.0), NEBNext Ultra Directional RNA Library Preparation Kit for Illumina (catalog number E7420, protocol revision 6.0), and NEBNext Multiplex Oligos for Illumina (catalog number E7600, protocol revision 2.0) with the following modifications: E7420, Section1.2 (“mRNA Isolation, Fragmentation and Priming Total RNA”) Step 37, we decreased the incubation time from 15 min to 5 min; E7420, Section 1.3 (“First Strand cDNA Synthesis”), Step 2, we increased the incubation time from 15 min to 50 min; for the size selection we used 40ul AMPure XP beads for the 1st bead selection and 20ul AMPure XP beads for the 2nd bead selection (targeting an insert size of 300-450bp and a final library size of 400-550bp); and in E7420, Section 1.9A (“PCR library Enrichment”), Step 2 we used 14 cycles for PCR cycling and dual index primers (i507 and i705-i712). Individual libraries were normalized to 10nM and eight samples were pooled per lane. Sequencing was performed at UC Berkeley’s QB3 Vincent J. Coates Genomics Sequencing Laboratory on an Illumina HiSeq4000 instrument, generating 150bp paired end reads.

### RNA-seq analysis

For each sample we produced 13-60M total reads and 4-30M mapped reads. Replicates were merged and aligned to the GRCh38.p8 version of the human genome using TopHat version: 2.1.1 (Trapnell et al. 2009; Kim et al. 2013). BAM files generated were used for the alternative splicing analysis and for the cassette reannotation. All the splicing-junction data formatted as bigBed and bigWig files obtained in this study were uploaded onto the UCSC genome browser; it can be accessed by copying the following link into a web browser: http://genome.ucsc.edu/cgi-bin/hgTracks?db=hg38&hubUrl=https://sina.lbl.gov/seqdata/conboy-ucsc/hub.txt. The FASTQ files were run through the STAR aligner (Dobin et al. 2013) to produce expression scores as input for DEXSeq (Anders et al. 1984; Reyes et al. 2013)

### Custom Cassette Reannotation Scripts

A custom reannotation tool was written using the Rust language to reannotate an input annotation to include additionally discovered cassette exons. The source code can be found at GitHub: https://github.com/bdgp/cassette_reannotation. The reannotation tool was given a curated subset of the NCBI RefSeq annotation version GCF_000001405.34 based on GRCh38.p8. This curated subset only included features beginning with NM, NR, and YP. TopHat version: 2.1.1 (Trapnell et al. 2009; Kim et al. 2013) generated alignment files of the sample reads were also passed in to the re-annotation tool. The tool finds constitutive splice pairs in the input annotation, then searches for overlapping reads in the input read alignment files that provide evidence for unannotated cassette exons. The tool requires at least two paired-end fragment splices for both the start and the stop of the cassette which also must splice at least one of the annotated flanking exons. The tool also requires contiguous read coverage throughout the discovered cassette. The tool then produces a re-annotation that includes all of the features of the input annotation, along with newly created transcript features. The newly created transcript features are based on transcript features existing in the input annotation, but with the discovered cassettes added.

### Alternative Splicing Analysis

The re-annotated annotation including discovered cassettes, along with the alignment files, were passed to SplAdder (Kahles et al. 2016) http://github.com/ratschlab/spladder to discover retained introns and alternative exons in the annotation. Some minor bug fixes, disabling of code assertions and parameter tuning were required for SplAdder to perform the analysis. We discovered that the least-stringent default parameter set was still to stringent to detect many retained introns in our sample data, so the individual parameters were made even less stringent. Details of the parameters used in our run of SplAdder can be found Supplementary Materials.

The retained intron adjusted PSI values returned by SplAdder unfortunately did not account for exons overlapping the retained intron, so a custom tool was written to correct the adjusted PSI values. SplAdder computes the adjusted PSI values using the average per-base coverage over the intron (intron_cov) and the number of reads with splices confirming the intron (intron_conf). The tool written to correct the adjusted PSI values recomputes the average per-base coverage by subtracting out the coverage of exonic regions. The tool uses this corrected intron_cov value along with the intron_conf directly from SplAdder to compute the corrected adjusted PSI value for the retained intron.

### Splicing reporter assays

A 4.7 kb region of the human SF3B1 gene extending from the 3’ end of intron 2 to the 5’ end of intron 6, was amplified using the following primers: F: 5’-tggaattctgcagatAAGGAGGGCTTAGACATCACAC-3’; R: 5’-gccagtgtgatggatCTATGGCAACCCAAGCAGA-3’. The fragment was cloned into pcDNA3.0 using In-Fusion methods (Gibson 2011) with 15nt in lower case sequence representing overlap with the ends of EcoRV-linearized vector. The splicing reporter was transfected into K562 using Fugene HD according to the manufacturer’s instructions (Promega). RNA was harvested after 48 hrs and purified according to the manufacturer’s instructions (Qiagen), but with the addition of a DNase step to eliminate potential contamination by genomic DNA. RNA was reverse transcribed with Superscript III (Invitrogen) into cDNA using the BGH reverse primer in the vector (5’-tagaaggcacagtcgagg-3’). Spliced products were amplified using a forward primer in exon 3 (5’-catcatctacgagtttgcttgg-3’) and a reverse primer in the vector (5’-atttaggtgacactatagaatagggc-3’). PCR reaction conditions were adjusted to allow for amplification of IR products ≥ 3 kb in length (denaturation at 95˚ C for 20sec, annealing at 56˚ C for 10sec, extension at 70˚ C for 2min 30sec; 35 cycles) using KOD polymerase in the presence of betaine to enhance amplification. PCR products were analyzed on either 2% agarose gels or 4.5% acrylamide gels. All PCR products discussed in the manuscript were confirmed by DNA sequencing. This strategy amplified minigene-derived transcripts but not endogenous SF3B1 mRNA, as confirmed using RNA from untransfected or empty vector-transfected cells.

### Enhanced CLIP analysis

Splicing maps were generated using U2AF1 (ENCSR862QCH) and U2AF2 (ENCSR893RAV) eCLIP normalized densities overlapped with selected retained introns containing candidate decoy exons at two time points (D9, D16). These density values were normalized first using an RPM transformation to account for variation in sequencing depth, then normalized again using equivalent densities from a size-matched input sample. To perform this second normalization, both IP and equivalent input signals were transformed into their probability densities to preserve overall shape of binding and to reduce signal dominance from a few events. Input densities for each event were then subtracted from the corresponding IP to remove background signal. The final density value represents the mean of these normalized densities devoid of any value exceeding the 95% median at each position to reduce confounding outlier effects.

In addition to candidate decoy exons, normalized densities were also overlapped with a set of cassette exons, derived from a subset of Gencode (v19) constitutive exons. Within these annotations, we define a ‘cassette’ as any exon found between 10% and 90% spliced in at least 50% of all shRNA knockdown control data (encodeproject.org, all non-specific target controls, aligned to hg19 with TopHat), filtering any region that is not supported by at least 30 reads. From this set, regions with the highest inclusion average between two replicates were chosen if any regions overlapped to remove any possibility of double counting eCLIP signal.

## ACKNOWLEDGEMENTS

This work was funded by NIH grant 5R01DK108020 (JGC) and by the Director, Office of Science and Office of Biological & Environmental Research of the US Department of Energy [DE-AC02-05CH1123]. BY and GWY are partially supported by the National Institute of Health under grants HG004659 and HG007005. We acknowledge Brenton Graveley’s laboratory for sharing RNA-seq datasets generated within the ENCODE project. This work used the Vincent J. Coates Genomics Sequencing Laboratory at UC Berkeley, supported by NIH S10 OD018174 Instrumentation Grant.

---

**Table 1.** Splicing reporters.

| Splicing reporter | Genome coordinates (GRCh38/hg38) |
| --- | --- |
| Wild type | chr2:197,416,658-197,421,257 |
| ΔE4ab | WT $\Delta$ chr2:197,420,152-197,420,358 |
| ΔE4c | WT $\Delta$ chr2:197,419,814-197,419,929 |
| ΔE4d | WT $\Delta$ chr2:197,419,575-197,419,677 |
| ΔE4e | WT $\Delta$ chr2:197,418,852-197,419,014 |
| ΔE4f | WT $\Delta$ chr2:197,418,719-197,418,815 |
| OGT | chrX:71,546,208-71,546,632 |
| ARGLU1 | chr13:106,557,456-106,557,720 |
| DDX39B | chr6:31,534,442-31,534,912 |
| SNRNP70 | chr19:49,101,950-49,102,379 |
| KEL | chr7:142,956,987-142,957,311 |
| FUS | chr16:31,187,786-31,188,063 |
| mut24.2 | E4e chr2:197,418,851-197,419,013<br>inserted in i5 after position chr2:197,417,934 |
| mut31 | 1.5kb of i4 sequences (chr2:197,418,852-197,420,358) inserted in place of<br>1.25 kb of i5 sequence (chr2:197,418,326-197,417,086) |
| Decoy exon regions from OGT, ARGLU1, DDX39B, SNRNP70, KEL, and FUS were inserted in place of the E4e region chr2:197,418,795-197,419,034 |  |

**Figure S1.**
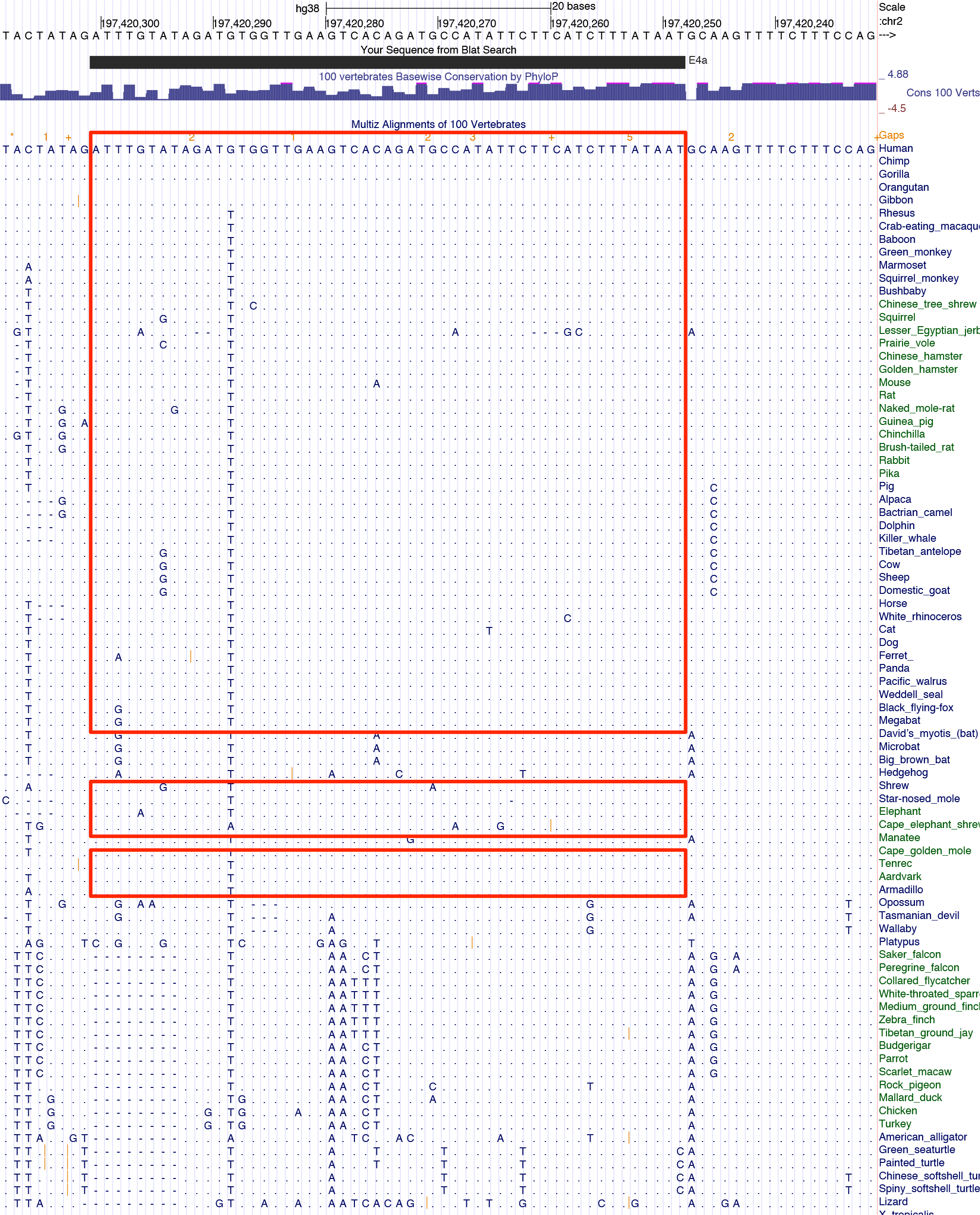
Conserva1on of SF3B1 exon 4a in most mammals.

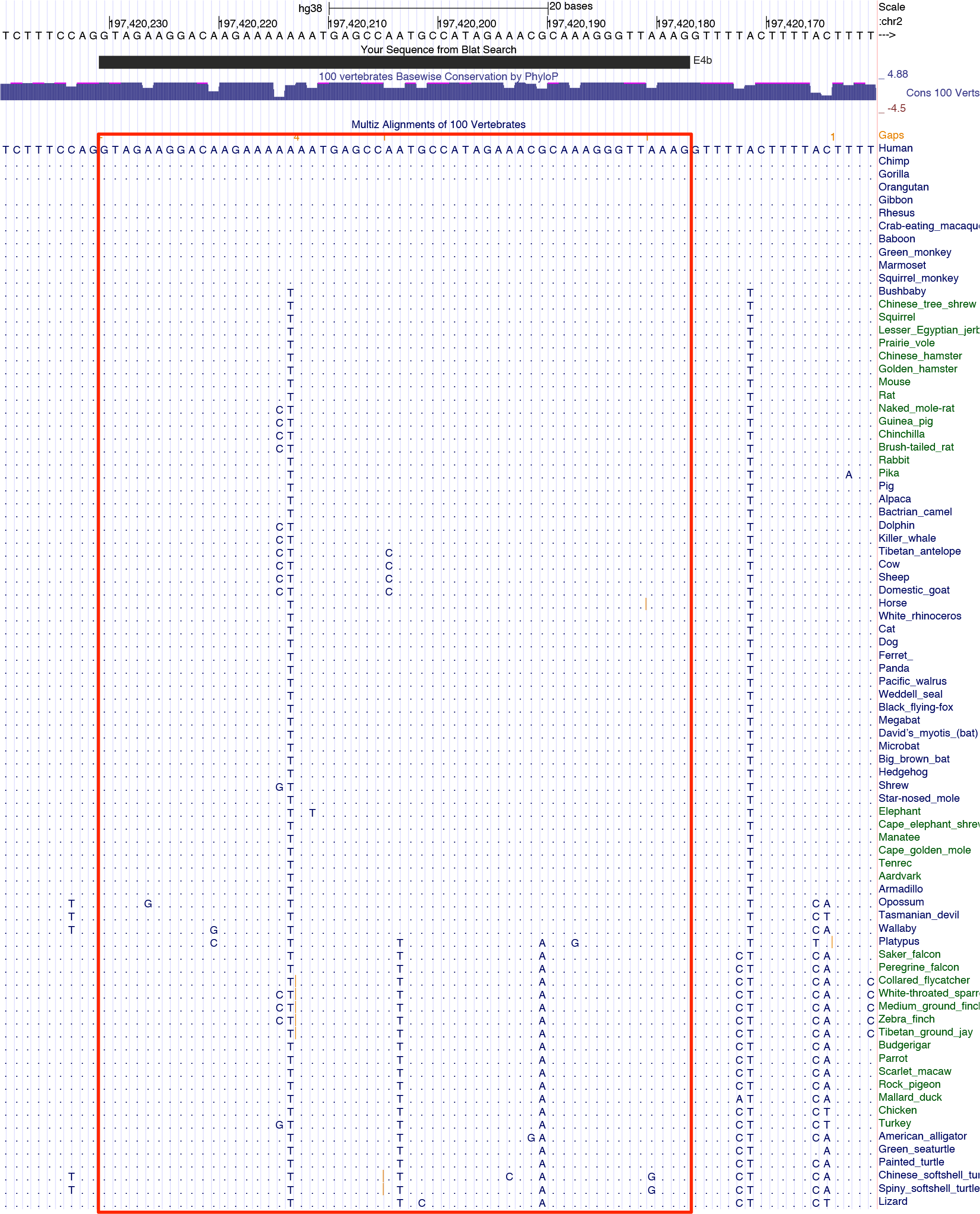
Conserva1on of SF3B1 exon 4b to repti1es.

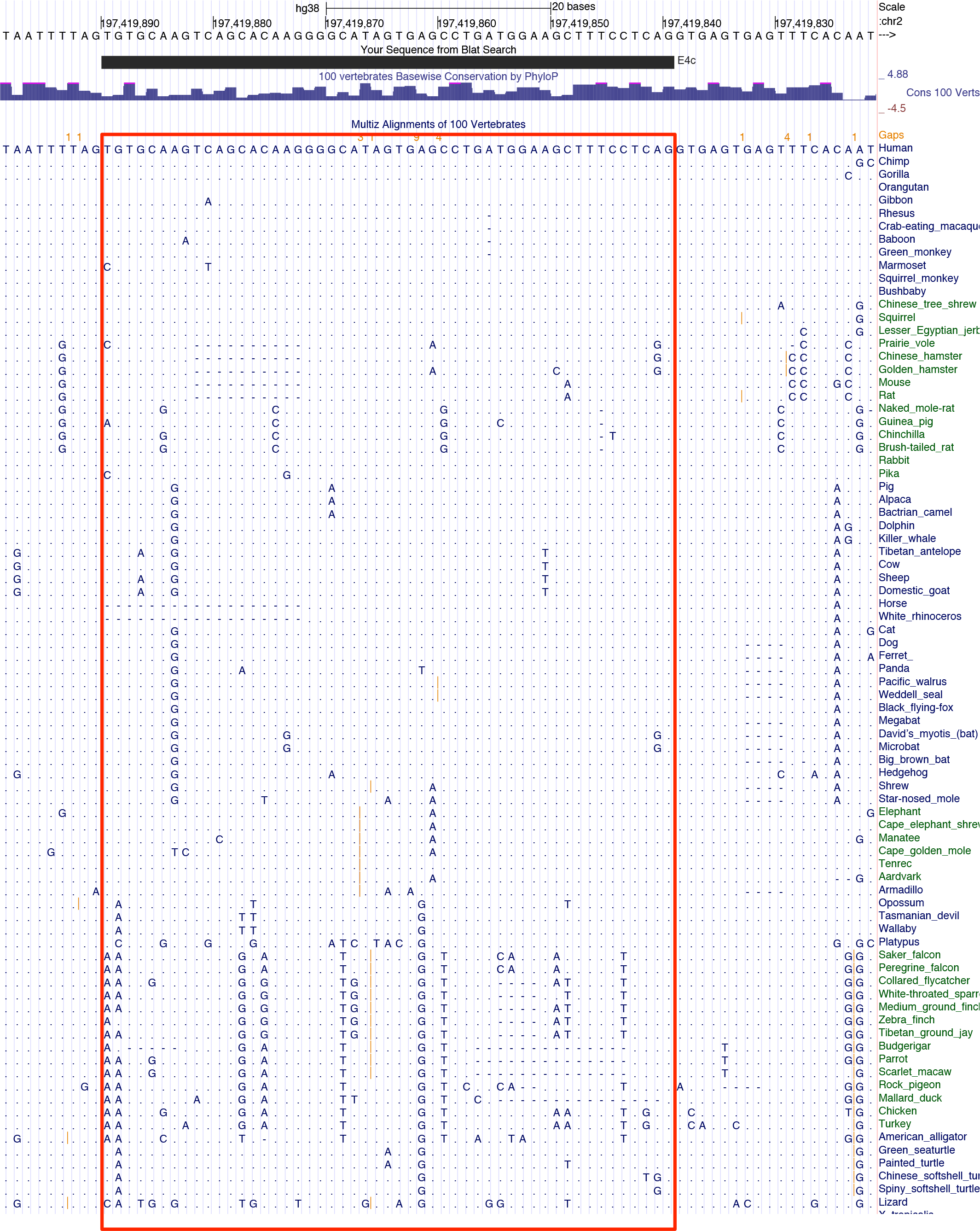
Conserva1on of SF3B1 exon 4b to repti1es.

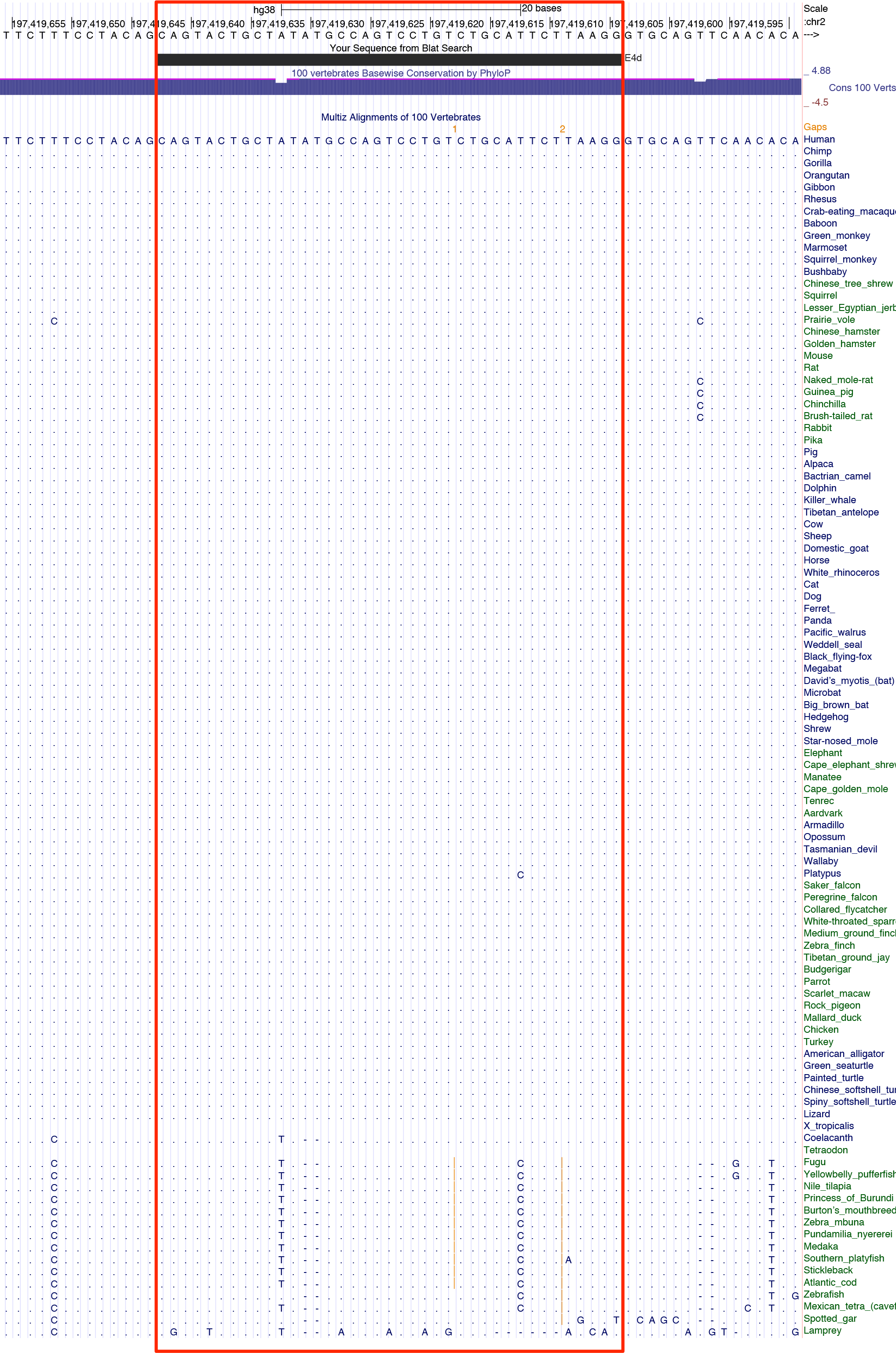
Conserva1on of SF3B1 exon 4d to fish genomes.

**Figure S1_5.**
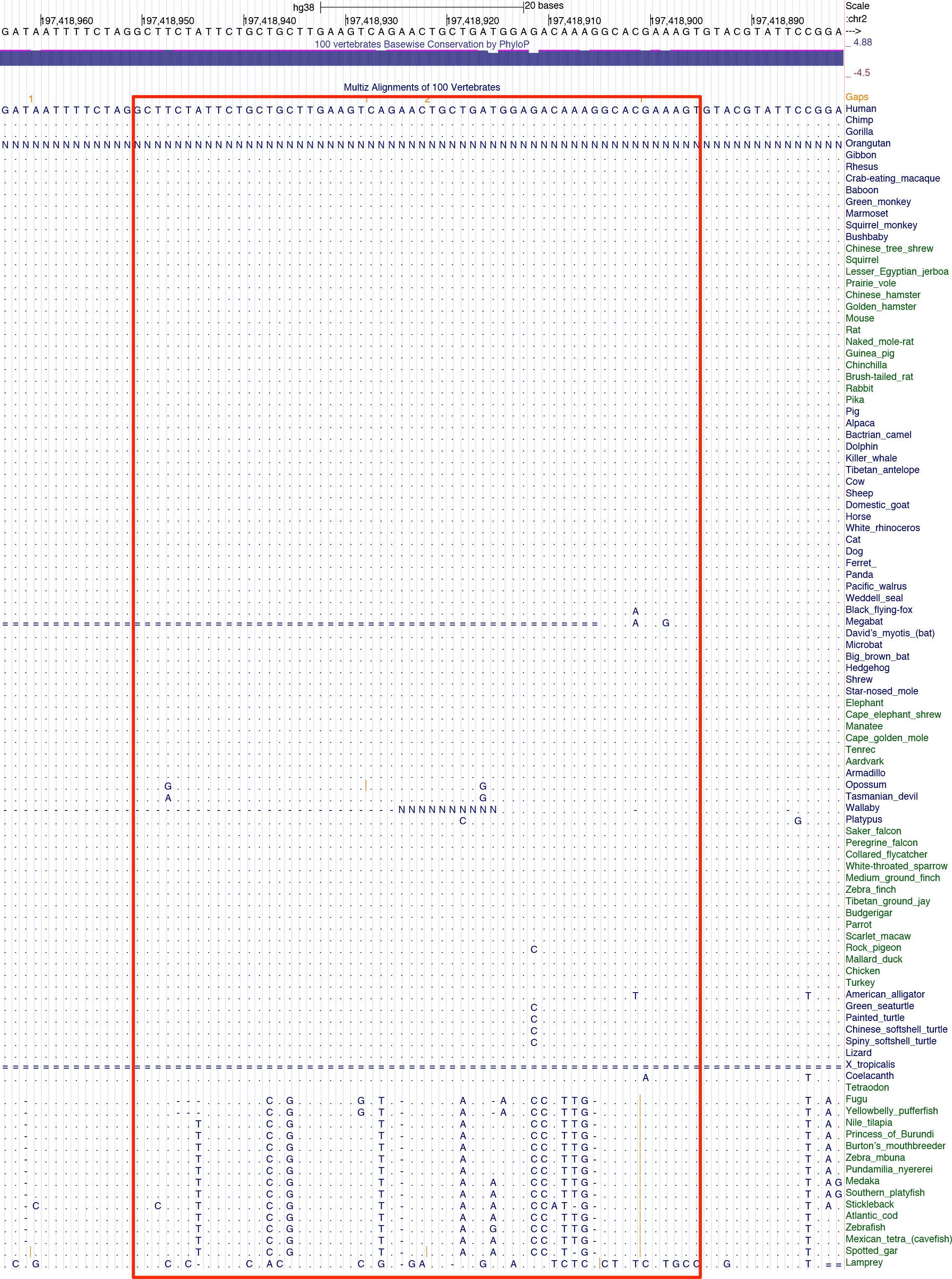
onserva1on of SF3B1 exon 4d to fish genomes.

**Figure S1_6.**
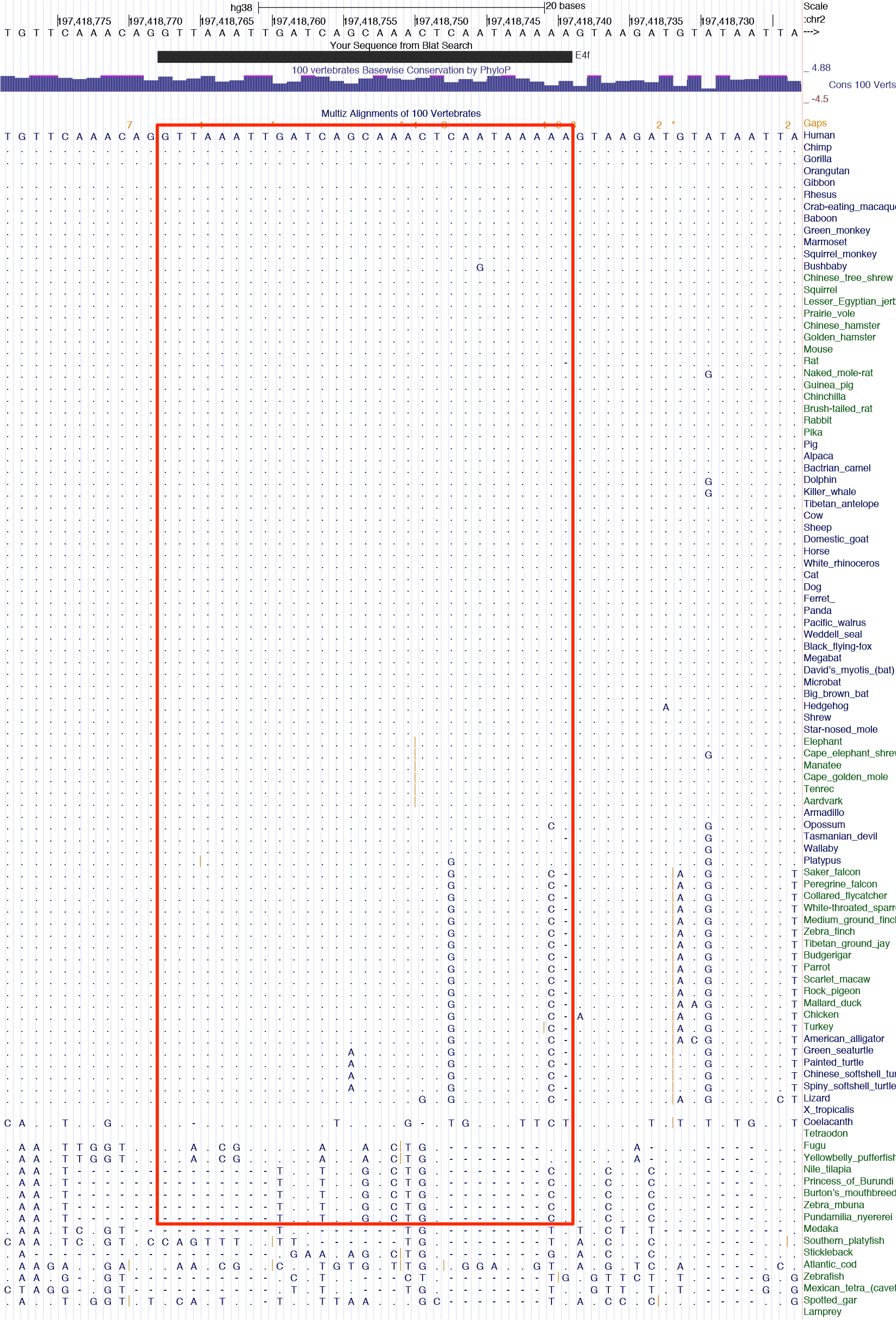
Conserva1on of SF3B1 exon 4e to fish genomes.

**Figure S2.**
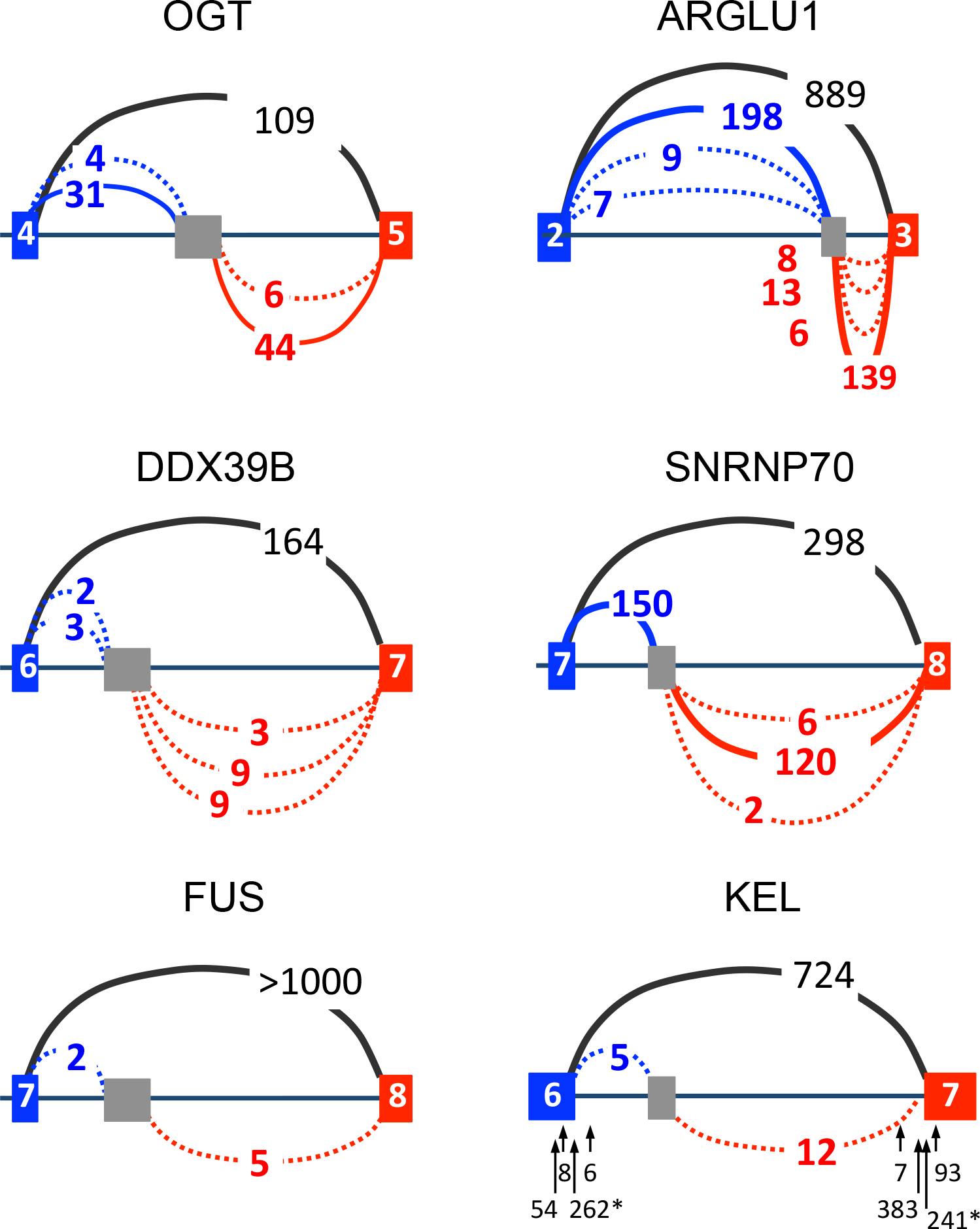
Splice junctions defining candidate decoy exons in selected erythroblast retained introns.

**Figure S3.**
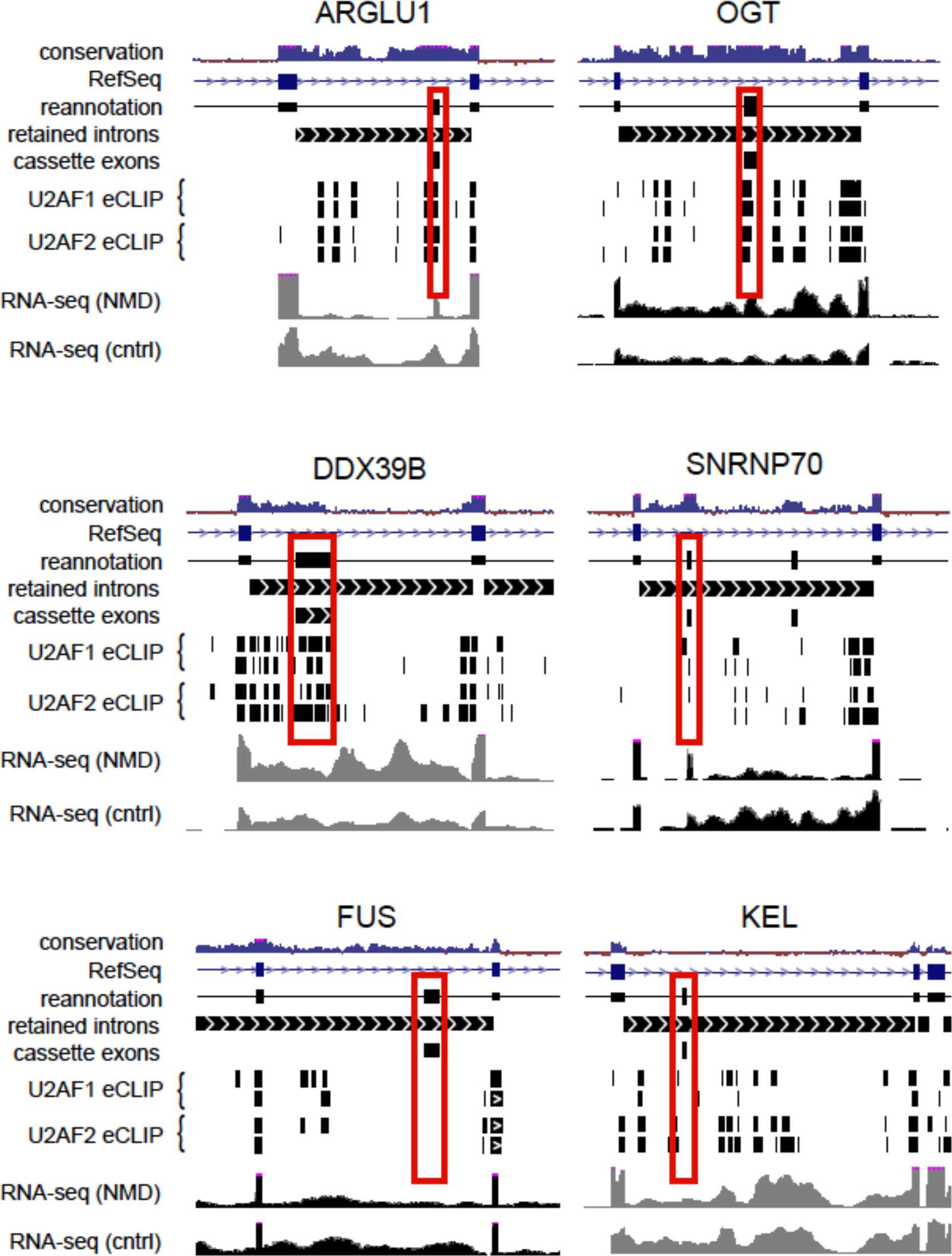
Genome browser tracks showing gene structure and expression feature of representative retained introns. Red boxes highlight predicted cassette exons that reside in retained introns. Considerable overlap with U2AF binding peaks was observed for some decoys (ARGLU1, OGT, DDX39B), but overlap was reduced (SNRNP70, KEL) or absent (FUS) in others.

